# Multi-omic analysis of maize NILs for chilling tolerance QTLs uncover regulatory and metabolic signatures

**DOI:** 10.64898/2026.03.04.709568

**Authors:** Maxence James, Andrea Rau, Anca Lucau-Danila, Jean-Michel Saliou, Bertrand Gakiere, Caroline Mauve, Alexandra Launay-Avon, Christine Paysant Le Roux, Stéphane Bernillon, Pierre Pétriacq, Catherine Giauffret, Estelle Goulas

**Affiliations:** Department of Molecular Sciences, Uppsala BioCenter, Swedish University of Agricultural Sciences and Linnean Center for Plant Biology, Uppsala, Sweden; Université Paris-Saclay, INRAE, AgroParisTech, GABI, F 78350 Jouy-en-Josas, France; Université de Lille, UMRT 1158 BioEcoAgro, Institut Charles Viollette, F 59000, Lille, France; Université de Lille, CNRS, INSERM, CHU Lille, Institut Pasteur de Lille, US41 - UAR2014 - PLBS, F 59000 Lille, France; Platform Métabolisme-Métabolome, Institute of Plant Sciences Paris-Saclay IPS2, CNRS, INRAE, Rue de Noetzlin, F 91192 Gif sur Yvette, France; Platform Transcriptomique POPS, IPS2, Institute of Plant Sciences - Paris-Saclay, Bâtiment 630, rue de Noetzlin, Plateau du Moulon, F 91190 - Gif-sur-Yvette, France; INRAE, RU1264 Mycology and Food Safety (MycSA), 71, Avenue Edouard Bourlaux, Villenave d’Ornon 33883, France; Univ. Bordeaux, INRAE, Biologie du Fruit et Pathologie, UMR 1332, F-33140 Villenave d’Ornon, France; AGAP Institut, Univ Montpellier, CIRAD, INRAE, Institut Agro, F34398 Montpellier, France; CNRS, UMR 8576 –Unité de Glycobiologie Structurale et Fonctionnelle, University of Lille, Lille, France; Bordeaux Metabolome, MetaboHUB, PHENOME-EMPHASIS, Villenave d’Ornon 33140, France

**Keywords:** Maize (*Zea mays*.L.), cold stress, chilling tolerance, genotype, near-isogenic lines, stress signaling, multi-omics, proteome, benzoxazinoids, cell wall remodeling, quantitative trait loci (QTL)

## Abstract

Early sowing of maize (Zea mays L.) is increasingly required to mitigate summer drought under climate change, making the acquisition of chilling tolerance a major agronomic challenge. Here, we investigated the molecular and physiological bases of cold tolerance using two maize near-isogenic lines (NILs) differing at two major chilling tolerance quantitative trait loci (QTLs) located on chromosome 4. Plants were exposed to low temperature (14°C day/10°C night) for 20 days and analyzed using an integrated multi-omics approach combining transcriptomics, soluble and cell wall proteomics, and metabolomics (primary and specialized metabolites), together with physiological measurements. Univariate and multivariate analyses revealed significant chilling-induced variability across all molecular layers, affecting ∼0.2% of genes, ∼6% of proteins, and a subset of specialized metabolites, while primary metabolites were largely stable. Integrative statistical analyses demonstrated that the soluble and cell wall proteomes contributed most strongly to the genotype effect, highlighting protein-level regulation as a major determinant of chilling tolerance. A restricted 5.15 Mb divergence region on chromosome 4 was sufficient to drive contrasting physiological responses, including differences in photosynthetic charge separation efficiency and leaf development, favoring the chilling-tolerant NIL. Notably, several components of the benzoxazinoid pathway located within the divergence region, including BX1 and associated specialized metabolites (BZX-like glucoside, DIBOA-glucoside-2, HBOA-glucoside-2), were specifically associated with chilling tolerance, suggesting a role in stress signaling and hormonal crosstalk. Overall, this study demonstrates that integrative multi-omics analyses provide a powerful framework to resolve genotype-specific regulatory mechanisms underlying chilling tolerance in maize and to identify candidate molecular targets for breeding.

**Highlights:**

- First organ-resolved multi-omics dissection of chilling responses in maize NILs.
- A 5.1Mb divergence on chromosome 4 drives major physiological and molecular differences.
- Chilling tolerance is linked to more robust photochemical homeostasis and sustained leaf development.
- Soluble and cell-wall proteomes dominate the genotype-discriminating -omics signal.
- Benzoxazinoids and defense-related transcriptional modules are differentially activated.
- Cell wall remodeling enzymes and apoplastic peroxidases emerge as key tolerance players.

## 1. Introduction

Plants are sessile organisms in a constantly changing environment and are therefore susceptible to various stresses. These include biotic stresses, such as pathogens, and abiotic stresses, such as nutrient deficiency temperature fluctuations. Abiotic stresses are currently defined as environmental conditions that can reduced the yield of a field crop species by up to 50% (Bray et al., 2000; Cramer et al., 2011; Kopecká et al., 2023). Among these conditions, temperature is a well-known factor limiting productivity and thus affecting the geographical distribution of species (Schepers et al., 2024; Zhu, 2016). To achieve optimal growth and then yield, plants require a daytime temperature range or an optimal temperature (Guy et al., 2008; Praat et al., 2024). Decreasing the temperature below the optimal value will result in a change in biochemical processes such as a decrease in the rate of metabolic reactions due to less active and unstable enzymes and an increase in membrane rigidity (Fonteyne et al., 2016; Marocco et al., 2005). However, plants can adapt their physiological processes to cope with moderate thermal stress (Jiang et al., 2025; Leipner and Stamp, 2009). Some plants are able to use this adaptation to further survive freezing events. It is then being called cold acclimation. But being a tropical species, *Zea mays* L., member of the *Poaceae* family, is strongly impacted by chilling stress (temperatures below 15°C) which becomes severe below 5°C (Riva-Roveda et al., 2016). Temperatures in the range of 10-15°C will result in decreased and delayed maize growth (Greaves, 1996). However, in recent decades, its cultivation in so-called cool climate regions has been made possible by an efficient breeding for chilling tolerance leading to better yields (Božić et al., 2024; Marocco et al., 2005; Riva-Roveda et al., 2016; Waititu et al., 2021). With climate change, strengthening chilling tolerance is especially relevant for enabling earlier sowing dates, which offer multiple agronomic benefits: the potential to cultivate longer-cycle, higher-yielding hybrids, avoidance of drought and heat stress during flowering, and a reduced risk of mycotoxin contamination caused by late-season *Fusarium* infections. Therefore, the issue of chilling tolerance is becoming more and more crucial and classical breeding approaches seem to have reached their own limits and require better knowledge of the mechanisms of chilling stress responses.

Cold stress rapidly activates an integrated signaling network in plants, in which phytohormones function as primary transducers linking temperature perception to transcriptional reprogramming (Eremina et al., 2016; Shi et al., 2015). Brassinosteroids and ABA are among the most influential regulators, respectively enhancing stress tolerance in chilling-sensitive crops and promoting the induction of CBF transcription factors (Knight et al., 2004; Xia et al., 2009). These early hormonal cues trigger a broad reshaping of gene expression across pathways associated with photosynthetic efficiency, secondary metabolism, and signal transduction (Kaplan et al., 2007; Li et al., 2019). Recent studies highlight a substantial contribution of jasmonate- and ethylene-dependent modules to cold acclimation in maize and other plants, suggesting a more complex hormonal architecture than previously assumed (Hewedy et al., 2023; Pérez-Llorca et al., 2023; Yang et al., 2023). Metabolite accumulation form also essential components of the chilling response (Urrutia et al., 2021). Primary metabolites such as amino acids, polyamines, and soluble sugars exert protective functions ranging from osmotic regulation to ROS buffering (Krasensky and Jonak, 2012). Proline is one of the most robustly induced metabolites in maize during chilling, reflecting its multifunctionality in maintaining cellular integrity (Guy et al., 2008; Moradtalab et al., 2018). Cold exposure also drives a reorientation of specialized metabolism. Benzoxazinoid biosynthesis genes, for instance, are strongly upregulated in chilling-treated maize, marking a shift in the chemical defense landscape (Li et al., 2017; Yu et al., 2021). Flavonoids also accumulate in parallel with declines in photosynthetic performance (Guy et al., 2008), and high-resolution metabolomics recently revealed increased flux through coordinated flavonoid metabolic reprogramming during early cold adaptation in tomato germplasm (Zhang et al., 2025). Structural remodeling of the cell wall constitutes another crucial layer of cold acclimation. Maize exposed to chilling typically shows elevated cellulose and reduced pectin content (Bilska-Kos et al., 2017; Sobkowiak et al., 2016), whereas cold acclimation leading to freezing tolerance is associated with increased pectin deposition and modification via pectin methyl esterase (Qu et al., 2011; Solecka et al., 2008). New cell-wall-omics analyses further implicate hemicellulose remodeling, particularly arabinoxylan branching, as a hallmark of the cold acclimated state (Domon et al., 2013; Takahashi et al., 2024). Overall, cold tolerance emerges from the coordinated action of hormonal cues, transcriptional regulators, metabolic pathways, and dynamic cell wall restructuring-highlighting the necessity of integrated multi-omics approaches to fully decode the complexity of chilling adaptation in maize.

Although transcriptomics has long been used to characterize plant responses to abiotic stress, it is now well established that RNA abundance alone rarely provides an adequate explanation for cold-induced physiological adjustments in maize. Transcript levels often correlate poorly with protein abundance (Chen et al., 2022; Renaut et al., 2006), and the extensive layers of translational, post-translational, and metabolic regulation further complicate interpretation when relying on a single molecular layer. The functional consequences of transcriptomic shifts are instead manifested across higher-order molecular layers-including the proteome and metabolome-whose regulatory dynamics are essential to understand the architecture of chilling tolerance. As a result, multi-omics approaches have become indispensable for deciphering the complexity of maize responses to abiotic stress (Niu et al., 2024; Yu et al., 2024; Zivy et al., 2015). Such frameworks provide a more complete perspective on how plants perceive, transduce, and respond to chilling stress by revealing regulatory modules, metabolic reprogramming, and genotype-specific adaptations that remain undetected in transcriptome-only datasets. Recent multi-omics studies in maize and other cereals have underscored the value of integrating transcriptome, proteome, and metabolome data to capture cold-induced functional shifts. Proteomic and metabolomic analyses frequently uncover chilling-responsive pathways that only partially overlap with transcriptomic predictions, demonstrating the unique and complementary information contributed by downstream molecular layers (Gu et al., 2023; Urrutia et al., 2021). Integrated analyses of maize seedlings exposed to low temperatures have identified key regulatory nodes and metabolic adjustments that cannot be resolved using a single-omics approach (Yu et al., 2024), while multi-omics network analyses have revealed organ- or age-specific differences in gene, protein, and hormone regulation under osmotic stress (Niu et al., 2024). Together, these studies highlight how multi-omics frameworks strengthen our capacity to dissect regulatory bottlenecks, identify robust biomarkers, and ultimately obtain a mechanistic understanding of plant stress acclimation.

To advance this multi-layered perspective, we undertook an in-depth physiological and multi-omics characterization of maize chilling responses using two near-isogenic lines (M1, chilling-tolerant; M2, chilling-sensitive) that differ only in a 5 Mb region of chromosome 4 containing two QTLs associated with chilling tolerance, which has been recently characterized (James et al., 2026). We implemented a long-term chilling treatment spanning 30 days, a duration rarely employed in maize research but particularly valuable for identifying stable molecular determinants of chilling tolerance. This experimental design enabled us to focus on treatment-specific genotype effects and to quantify respective contributions of transcripts, proteins, and metabolites. The integration of phenotypic variation with multi-omics datasets allowed us to identify key molecular, biochemical, and metabolic players that shape the maize chilling response. These findings provide a comprehensive resource for understanding cold tolerance mechanisms and support the development of maize cultivars better adapted to the climatic challenges of the coming decades.

## 2. Materials and methods

### 2.1. Experimental procedures

Plant growth and harvests-Maize seeds from two different genotypes (chilling-tolerant M1 and chilling-sensitive M2) were germinated in Petri boxes between two pieces of wet blotting paper at 20°C in darkness for 4 days. The seedlings were then transplanted into pots filled with Klasmann TS3 potting soil. Plants were grown under growth chamber conditions until the 3-leaf stage. Photosynthetic photon flux density was 500 µmol m–2 s–1, and day and night temperatures were set to 25 and 22°C, respectively, for the control temperature treatment, and 16 and 14°C, respectively, for the chilling temperature treatment. Conditions henceforth refer to the combination of genotype (M1 and M2) and treatment (control and chilling). Plants were harvested when showing 4 fully expanded leaves, with the third leaf harvested and divided into two parts with respect to the median leaf vein. A total of 12 pooled samples (minimum of 30 plants per sample) corresponding to 3 biological replicates for the 2 genotypes (M1 and M2) submitted to the 2 treatments (control and chilling) were collected and stored at -80 °C until further -omics analyses.

Physiological measurements-Three representative plants for each condition were chosen for measurement of morphological and photosynthetic parameters. For morphological parameters, we measured the number of visible leaf-ligules and tips, shoot dry weight, length and width of leaf n°3. Concerning photosynthetic parameters, we analyzed chlorophyll content with CCM-200 (Opti-Sciences, Hudson, NH) chlorophyll-meter and maximum quantum yield (Fv/Fm) and PSII quantum yield in light of 500 μmol m–2 s–1 (φ PSII) with mini-PAM fluorimeter (Effeltrich, Germany).

### 2.2. Chemical analyses

Total RNA extraction and sequencing is described in James et al. (2026). Briefly, total RNAs of the twelve samples were extracted for each biological replicate from 100 mg of shoot samples using NucleoSpin® RNA Plant kit (#740949, Macherey Nagel, Düren, Germany) and purified using RNA Clean & Concentrator-5 kit (#ZR1013, Zymo Research, California, USA). Sequencing was performed on an Illumina NexSeq500 (IPS2, POPS platform). RNA-seq libraries were generated with the TruSeq_Stranded_mRNA_SamplePrep_Guide_15031047_D protocol (Illumina®, California, U.S.A.), using paired-end (PE) and stranded sequencing with size 260 bp and read length of 75 bases, random lane assignment, and barcoding, yielding approximately 15 million PE reads per sample. A total of 44146 transcripts were sequenced (Supplemental dataset S1).

#### 2.2.1. Primary metabolite extraction and detection

After extraction from the twelve samples, the primary metabolites were analyzed from 5 mg of lyophilized material. The ground dried samples were resuspended in 1 mL of frozen (-20°C) water:acetonitrile:isopropanol (2:3:3) containing Ribitol at 4 µg.mL-1 and extracted for 10 min at 4°C with shaking at 1500 rpm in an Eppendorf Thermomixer. Insoluble material was removed by centrifugation at 16 300 g for 10 min. In total, 100 µl were collected and 10 µL of myristic acid d27 at 30 µg mL-1 were added as an internal standard for retention time locking. Extracts were dried for 4 h at 35°C in a Speed-Vac and stored at -80°C. All steps for GC-MS analyses were carried out as previously described (Fiehn, 2006; Fiehn et al., 2008). Samples were taken out of -80°C, warmed to 15 min before opening and dried again in a Speed-Vac evaporator for 1.5 h at 35°C before adding 10 µL of 20 mg.mL-1 methoxyamine in pyridine to the samples; the reaction was performed for 90 min at 30°C under continuous shaking in an Eppendorf thermomixer. Here 90 mL N-methyl-N-trimethylsilyl-trifluoroacetamide (MSTFA) (Regis Technologies, Morton Grove, IL, USA) were then added and the reaction continued for 30 min at 37°C. After cooling, all samples were transferred to an Agilent vial for injection. At 4 h after derivatization, 1 µL of sample was injected in splitless mode on an Agilent 7890B gas chromatograph coupled to an Agilent 5977A mass spectrometer. The column was an Rxi-5SilMS from Restek (30 m with 10 m Integra-Guard column). An injection in split mode with a ratio of 1:30 was systematically performed for saturated compound quantification. Oven temperature ramp was 60°C for 1 min then 10°C min-1 to 325°C for 10 min. Helium constant flow was 1.1 mL.min-1. Temperatures were the following: injector: 250°C, transfer line: 290°C, source: 230°C and quadropole 150°C. The quadrupole mass spectrometer was switched on after a 5.90 min solvent delay time, scanning from 50 to 600 m/z. Absolute retention times were locked to the internal standard d27-myristic acid using the RTL system provided in Agilent’s Masshunter software. Retention time locking reduces run-to-run retention time variation. Samples were randomized. A fatty acid methyl esters mix (C8, C9, C10, C12, C14, C16, C18, C20, C22, C24, C26, C28, C30) was injected at the beginning of the analysis for external RI calibration. The Agilent Fiehn GC/MS Metabolomics RTL Library (version June 2008) was employed for metabolite identification. Peak areas were determined with the Masshunter Quantitative Analysis (Agilent Technologies, Santa Clara, CA, USA) in splitless and split 30 modes. Resulting areas were compiled into one single MS Excel file for comparison. Peak areas were normalized to Ribitol and Dry Weight. A total of 117 identified metabolites are thus expressed in arbitrary units (semi-quantitative determination; Supplemental dataset S4). The annotation of metabolites was performed by MetaboAnalyst, allowing various accession numbers such as ID_Pubchem, ID_HMDB to be obtained.

#### 2.2.2. Specialized metabolite extraction and detection

The extraction of specialized metabolites by LC-MS analysis was adapted from Musseau et al., (2020). Briefly, 10 mg of freeze-dried leaf material were extracted with 1 mL methanol / water (70/30) containing 0.1% of formic acid and methyl vanillate as an internal control. Extracts were centrifuged and filtered before LC-MS injection. Blank extracts and quality control (QC) mix, which consisted of the mixture of the same volume of each sample extract, were also prepared. Methanolic extracts were injected on a LC-MS system including an Ultimate 3000 HPLC and an LTQ-Orbitrap Elite mass spectrometer (Thermo Fisher Scientific, Bremen, Germany) equipped with a HESI-II probe. Separation was performed with a C18 column (C18-Gemini, 2.0 x 150 mm, 3 µm, 110Å; Phenomenex, Torrance, CA, USA). Acquisitions were carried out in positive mode with the following parameters: spray voltage: 3.2 kV, sheath gas: 40 a.u., aux gas: 20 a.u., sweep gas: 3 a.u., capillary temp: 300 °C, source heater temp: 300°C, m/z range: 50-1500, resolving power 60k @ m/z 200. MS/MS spectra were acquired following a data-dependent acquisition strategy with a HCD mode and a normalized collision energy of 35%. Injections were randomized and QC mix was injected every 10 samples to correct for MS detector drift. Raw data were processed using the MS-DIAL v4.60 software (Tsugawa et al., 2015). Variables detected in blank extracts were filtered out. Intensity drift was corrected using LOWESS regression, retention time correction was performed using known metabolites, and zero-valued intensities were replaced by 10% of the minimum height. Finally, intensities were normalized according to the sample powder mass used for extraction. Finally, a total of 3569 variables were used for statistical analysis (Supplemental dataset S5) Annotation of benzoxazinoids was performed with retention time, accurate m/z, and MS/MS spectra using previous work (Urrutia et al., 2021).

#### 2.2.3. Soluble and cell wall protein enrichment

Soluble proteins (SP) were extracted from 2 g of frozen fresh leaves after grounding for 2 min in 10 ml Tris-HCl buffer (50 mM, 0.06% PIC (w/v, Protease inhibitor mixture), pH 7.5; Sigma-Aldrich) and then centrifuged (10 min at 4°C, 16,000g) as described in Goulas et al., (2001). The pellet for further cell wall protein (CWP) enrichment was frozen at -80°C until use, while the supernatant containing soluble proteins (SP) was recovered and quantified by the protein staining method (Bradford, 1976) using BSA. The supernatant was precipitated by incubation with 72% TCA (w/v, trichloroacetic acid, 2 h, 4°C), centrifuged for 10 min at 4°C and 16,000g, and the resulting pellet containing soluble proteins was washed once with cold acetone, dried at room temperature, resuspended in rehydration buffer (7 M Urea, 2 M Thiourea, 100 mM DTT, 2% w/v CHAPS) and stored at -80°C until use. CWPs enrichment was performed from the CWP pellet according to Printz et al., (2015). Briefly, the CWP pellet was suspended with CaCl2 buffer (5 mM NaAc, 200 mM CaCl2, pH 4.6) and incubated twice for 30 min at 4°C under gentle agitation. After each centrifugation (15 min at 4°C and 10,000g), the supernatant containing the CWPs was recovered and pooled. Then the solid cell wall residue was incubated twice with EGTA buffer (5 mM NaAc, 50 mM EGTA, pH 4.6) for 1 h at 37°C. After each centrifugation (15 min at 4°C and 10,000g), the supernatants containing the CWPs were recovered and polled. At last, the pellet was incubated with LiCl buffer (5 mM NaAc, 3 M LiCl, pH 4) overnight at 4°C with gentle agitation. The resulting supernatant after centrifugation (15 min at 4°C and 10,000g) was pooled with all previous supernatants resulting from the previous CWP steps to constitute the CWP extract. The CWP extract was concentrated using Amicon Ultra-15 3K Centrifugal Filter Device (Millipore) as recommended by the manufacturer, and CWP concentration was assessed by protein staining (Bradford 1976). The CWPs were then precipitated and resuspended in rehydration buffer (7 M Urea, 2 M Thiourea, 100 mM DTT, 2% w/v CHAPS) and stored at -80°C until further use.

#### 2.2.4. Soluble and cell wall protein detection

SP and CWP extracts were boiled for 10 min to denature the proteins, then cooled until required for SDS-PAGE as described by Laemmli, (1970) using a 5.5 % stacking gel and a 10 % separation gel in a Mini-PROTEAN® Tetra Cell Electrophoresis system. Equal amounts of protein were loaded per lane, according to the protein quantification results. Gels were stained with Bio-Safe Coomassie Stain (Bio-Rad) and the protein bands were excised for further protein identification and quantification by the XIC method. The first steps of discoloration / reduction / alkylation / dehydration / digestion / extraction of the peptides were performed using an automated MicroLAB NIMBUS system (Hamilton, Switzerland). The NanoLC-MSMS peptide analyses were then carried out on UltiMate 3000 liquid chromatography directly coupled with an Orbitrap Q-Exactive mass spectrometer (Thermo Scientific, USA). Raw data were processed using the Proteome Discoverer software (Thermo Scientific, USA), and identification was performed by the Mascot search engine (version 2.4.0, Matrix Science, London, UK) against a library composed of *Zea mays* L. protein sequences (Taxonomy 4577, Uniprot), conventional laboratory and technical contaminants, fused with the reverse sequences to identify proteins with a false positive rate of less than 1%. Only proteins present in all biological repetitions independently run by MS analysis were retained for further study. Proteins with missing values in only a subset of replicates for a given condition were imputed using K-nearest neighbors using the impute package with default parameters. Functional classification was performed using Panther (http://mapman.gabipd.org/web/guest/mapmanstore). Subcellular locations were predicted using WoLF PSORT (https://wolfpsort.hgc.jp/), TargetP1.1 (https://www.cbs.dtu.dk/services/TargetP/), and SignalP 4.1. (https://www.cbs.dtu.dk/services/SignalP/). The predicted cell wall location of proteins was verified using the WallProt database (https://www.polebio.lrsv.ups-tlse.fr/WallProtDB/), and proteins were functionally classified according to classes. If one protein was identified both in soluble and cell wall extracts, its final assignment was determined accordingly to its *in silico* localization predictions within XXX databases (i.e. proteins detected in an inconsistent fraction were reassigned accordingly). After this expert curration, a total of 5259 SPs and 201 CWPs were obtained (Supplemental dataset S2 and S3).

### 2.3. Bioinformatics and statistical analyses

The differential analyses were based on a contrast highlighting the treatment-specific genotype effect, referred to as the ‘**genotype effect’** where M1 (chilling-tolerant) was compared to the M2 (chilling-sensitive) genotype under chilling treatment and under control condition, i.e. Chilling (M1 vs M2) (Figure 1).

**Fig. 1.**
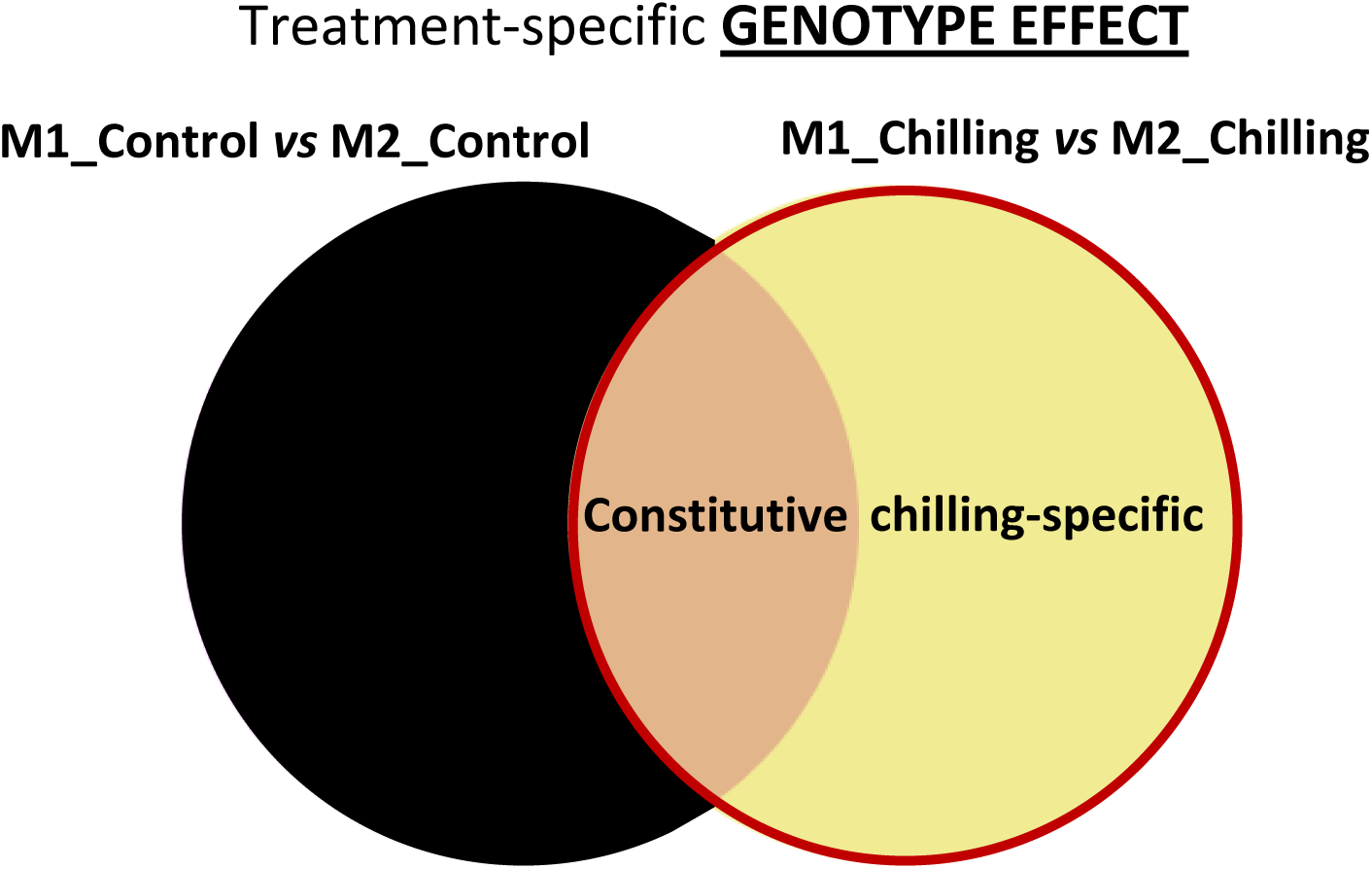
Venn diagrams of the two contrasts applied for statistical differential analyzes with -omics datasets A) genotype-specific treatment effect, also called “treatment effect” (M1+M2 (Control *vs* Chilling)) and B) constitutive- and chilling-specific genotype effects, also called “genotype effect” (Chilling (M1 *vs* M2)).

#### 2.3.1. RNA-seq analysis

RNA-seq analysis is described in James et al., (2026). Briefly, contrasts were defined to perform comparisons of interest, using Benjamini-Hochberg (BH) control of false discovery rate (FDR) to adjust for multiple testing. Only genes with absolute log_2_ fold change ≥1 and an adjusted p-value < 0.05 were considered to be differentially expressed genes (DEG) (Sekhon et al., 2019). Uniprot accession number, protein sequences and Gene Ontology (GO) annotations were retrieved for all DEGs with the uniprot database (https://www.uniprot.org/). An extensive functional characterization of DEGs was performed using gene ontology (GO) annotations and Family/Subfamily with the Panther database (http://www.pantherdb.org/, Mi et al., 2019).

#### 2.3.2. Proteomic analysis

Differential analyses were performed on log_2_ standardized imputed data with a linear model including fixed effect for genotype, using the limma Bioconductor package (V3.42.2, Ritchie et al., 2015). As for the transcriptome data, contrasts were used to test the significance of comparisons of interest, and p-values were adjusted with the BH method. Proteins with absolute fold changes ≥1.5 and adjusted p-value < 0.05 were considered to be differentially abundant.

#### 2.3.3. Metabolomic analysis

Differential analyses were performed on standardized data as described above for the proteomics data. Regardless of the value of log_2_ fold-change, metabolites with an adjusted p-value < 0.05 were considered to be differentially abundant (Kang et al., 2019).

#### 2.3.4. Multi-omics analyses

Differences in the expression of genes or abundance of proteins and metabolites were identified using the aforementioned differential analyses to compare chilling and control treatment or M1 and M2 genotypes. To visualize global structuring related to treatment or genotype effects, principal component analyses (PCA) were performed on normalized data within each -omics dataset individually with the R-package mixOmics (V6.10.6, Rohart et al., 2017) and ropls R packages (V1.18.8, Thevenot et al., 2015). Multiple factor analysis (MFA), a multi-table extension of PCA, was also performed to explore multi-omic data structures across the transcriptome, proteome, and metabolome using the FactomineR R package (V2.4).

#### 2.3.5. Enrichment analyses

GO enrichment analyses were performed for the Biological Process (BP) ontology for lists of genes or proteins with at least 100 elements using the topGO Bioconductor package (V2.54.0). A matched background set of the 10 closest genes for each element in the list (according to the Manhattan distance with respect to mean normalized expression) was used as the reference universe. Results focused on terms identified with a p-value < 0.05 according to the “weight01” algorithm and Fisher test statistic and at least 2 significant genes. A semantic classification of identified terms was performed with the GOGO package (Zhao and Wang, 2018).

## 3. Results

### 3.1. Morphological and photosynthetic features

Regarding the total number of visible leaves (Figure 2A), no differences were noticed between the chilling-tolerant genotype M1 and the chilling-sensitive genotype M2 under control conditions. After chilling, both genotypes showed a reduced number of total leaves, which was then different between M1 and M2 genotypes. No impact of the temperature on the number of ligulated leaves could be observed for a given genotype, this number being always higher in M1 than in M2 genotype. The shoot dry weights, the leaf lengths (Figures 2D and 2E) were also always higher in control conditions compared to chilling conditions, both for the chilling-tolerant genotype M1 and the chilling-sensitive genotype M2. The shoot dry weight was especially affected by chilling conditions with a decrease of 77% in M1 and 83.8% in M2. Whatever the temperature regime, control or chilling, no significant differences could be observed between M1 and M2 genotype neither for shoot dry weight, nor for leaf length and leaf width (Figures 2D, 2E and 2F). All three photosynthetic parameters, chlorophyll contents, maximum quantum yields (Fv/Fm) and PSII quantum yields (φ PSII) were significantly drastically lowered under chilling condition regarding to the control condition (Figure 2G, 2H and 2I), whatever the genotype considered. The only difference between the 2 genotypes within one condition was observed for the maximum quantum yield (Fv/Fm) under chilling conditions, with a decrease of nearly 20% in M2 compared to M1 (Figure 2H).

**Fig. 2.**
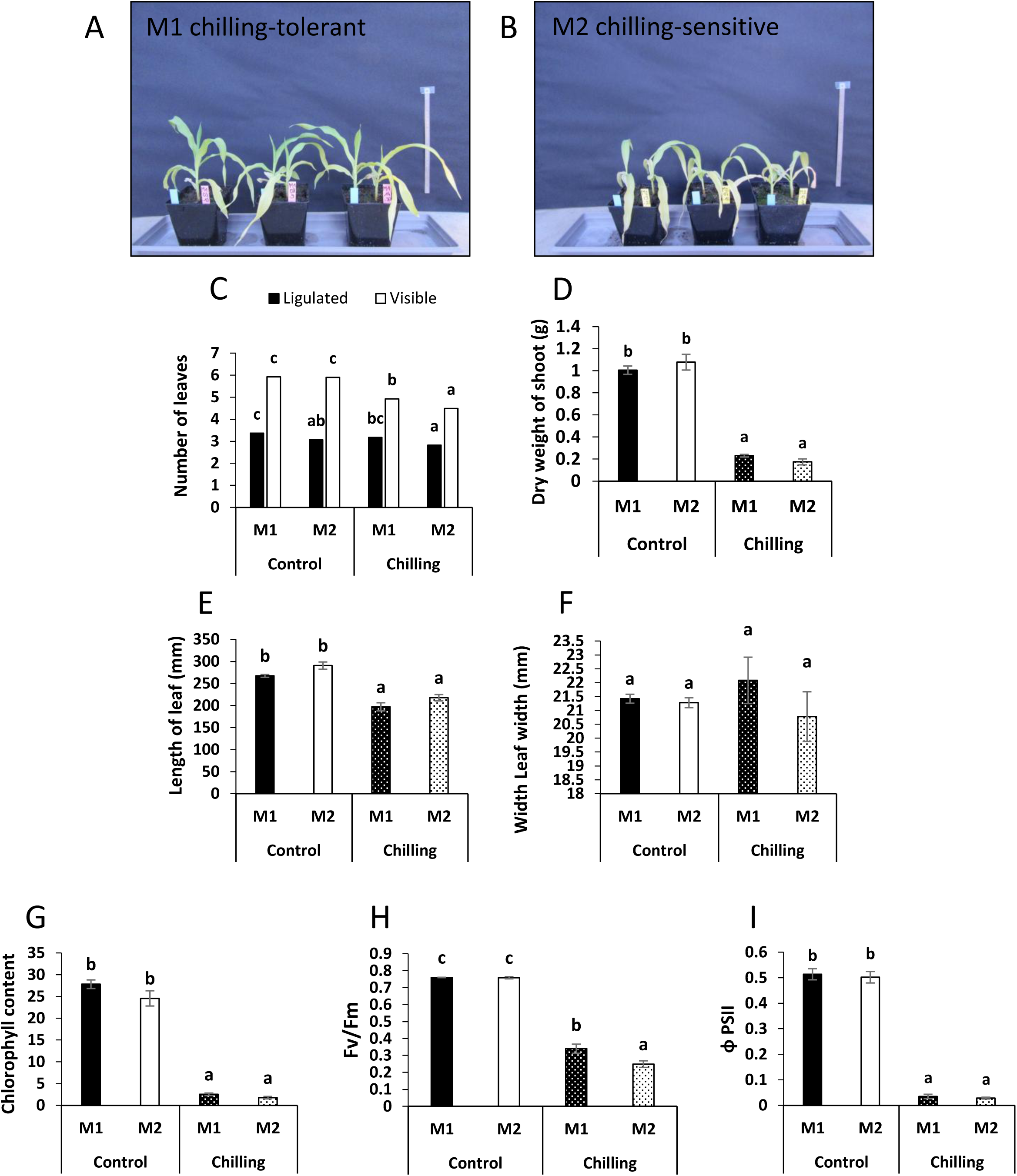
Morphological and photosynthetic parameters at 25/22 °C and 16/14 °C in M1 chilling-tolerant and M2 chilling-sensitive genotypes. Phenotypic appearance of genotypes A) M1 and B) M2 during chilling treatment. Morphological parameters correspond to C) the number of ligulated and visible leaves, D) the dry weight of shoot and E) the length and F) the width of leaf n°3. Photosynthetic parameters correspond to G) the chlorophyll content, H) the maximum quantum yield (Fv/Fm) and I) the PSII quantum yield in light of 500 μmol.m–2.s–1 (φPSII).*The analyzes were performed on maize leaves of M1 chillling-tolerant and M2 chilling-sensitive genotypes subjected either to control or chilling treatment. Results are presented as means ± SE (n=3). Significant differences between genotype and thermal condition are indicated by different letters (p-value≤0.05)*.

### 3.2. Multivariate analysis of -omics

Principal component analyses (PCA) were performed on each -omics datasets to identify global structures specifically related to the chilling treatment or genotype (Figure 3A-E). Across omics, the first principal component (PC1) was found to be strongly related to the treatment effect. PC1 explained 54%, 36%, 52%, 61% and 68% of the total variability for the transcriptome, soluble and cell-wall proteome, primary and specialized metabolome datasets, respectively (Figure 3A, 3B, 3C, 3D and 3E). PC2 instead appeared to be related to a genotype effect for the soluble and cell-wall proteome, explaining 20% and 15% of the total variability, respectively (Figure 3B, 3C); this structure was not so obvious in PC2 for the remaining omics. A Multiple Factor Analysis (MFA) (Figure 3F, 3G) performed on the transcriptome, soluble proteome, cell-wall proteome, primary metabolome and specialized metabolome datasets showed a clear separation on the first dimension (52% of the total variability) between samples subjected to the chilling and control treatments (Figure 3F). Variability on the second MFA axis (11% of the total variability) instead appears to be related to the two genotypes, albeit to a lesser degree. All five -omics datasets have similar coordinates on the first axis (Figure 3G), suggesting that this first MFA component related to the treatment effect is largely shared among them. However, the soluble proteome data (∼60%), and to a smaller amount the cell wall proteome data (∼25%), have the largest coordinates on the second axis, implying that they have a stronger contribution to this genotype-related component.

**Fig. 3.**
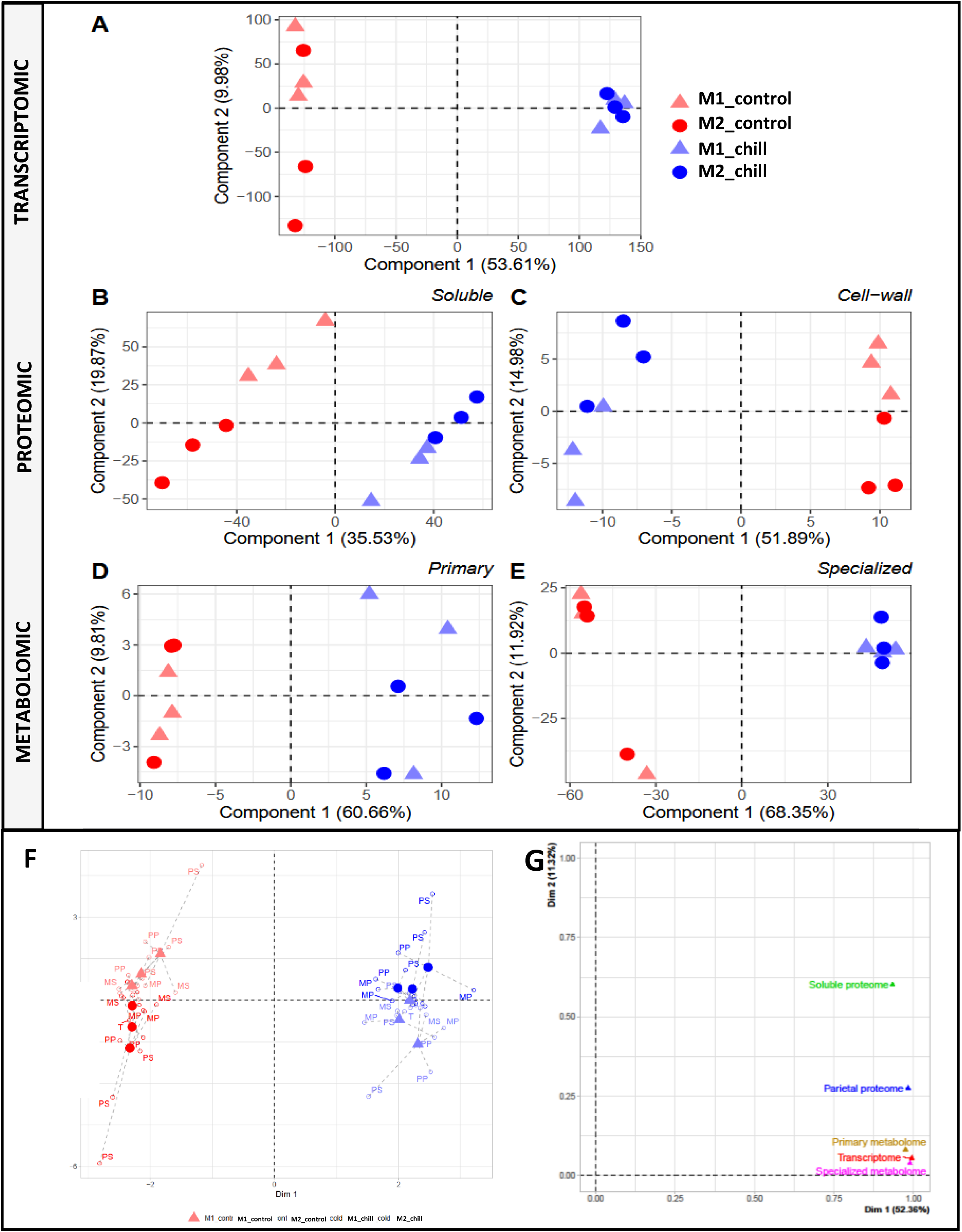
Principal Component Analysis (PCA) score plots of A) gene expression, B) soluble protein-, C) cell-wall protein-, D) primary metabolite-E) specialized metabolite contents. Multiple factorial analysis (MFA) score plots performed by FactoMineR package of R software showing the impact of the transcriptome, soluble and cell-wall proteome, primary and specialized metabolome on the variability of F) each different samples and G) –omics groups. A*nalyzes were based on differentially expressed or abundant gene, proteins and metabolites profiling datasets of maize leaves of M1 chilling-resistant and M2 chilling-sensitive genotypes submitted either to control (red) or chilling treatment (blue)*.

### 3.3. RNA-seq

A total of 55 DEGs were observed including 26 up-regulated and 29 down-regulated (Figure 4; Table S5 and S6). DEGs were represented by their functional annotation with a histogram (Figure 5), which indicated many biological functions including response to biotic and abiotic stress, transcription and translation regulation, carbon fixation / photosynthesis, cell wall organization and protein folding. 17 genes, representing approximately 31% of the DEGs, played a role in the response to biotic and abiotic stress, of which 7 DEGs were up-regulated and 10 down-regulated in the M1 chilling-tolerant genotype compared to the M2 sensitive-genotype. Genes involved in the regulation of transcription and translation were particularly up-regulated in the M1 genotype, with a total of 4 genes *versus* 1 down-regulated. Carbon fixation and photosynthesis functions were strongly impacted with 2 up-regulated genes but especially 4 down-regulated genes in the resistant genotype. There were many genes specifically up-regulated in the resistant M1 genotype compared to the sensitive M2 genotype, with functions in signaling, growth and development and mitochondrial biogenesis with for example 2 DEG with a function in signaling. In addition, specifically down-regulated genes in the M1 genotype were involved in transport, energy metabolism, ribosome maturation, lipid metabolic process. A large group of up- and down-regulated genes, representing 24% of DEGs with the genotype effect, had unknown functions.

**Fig. 4.**
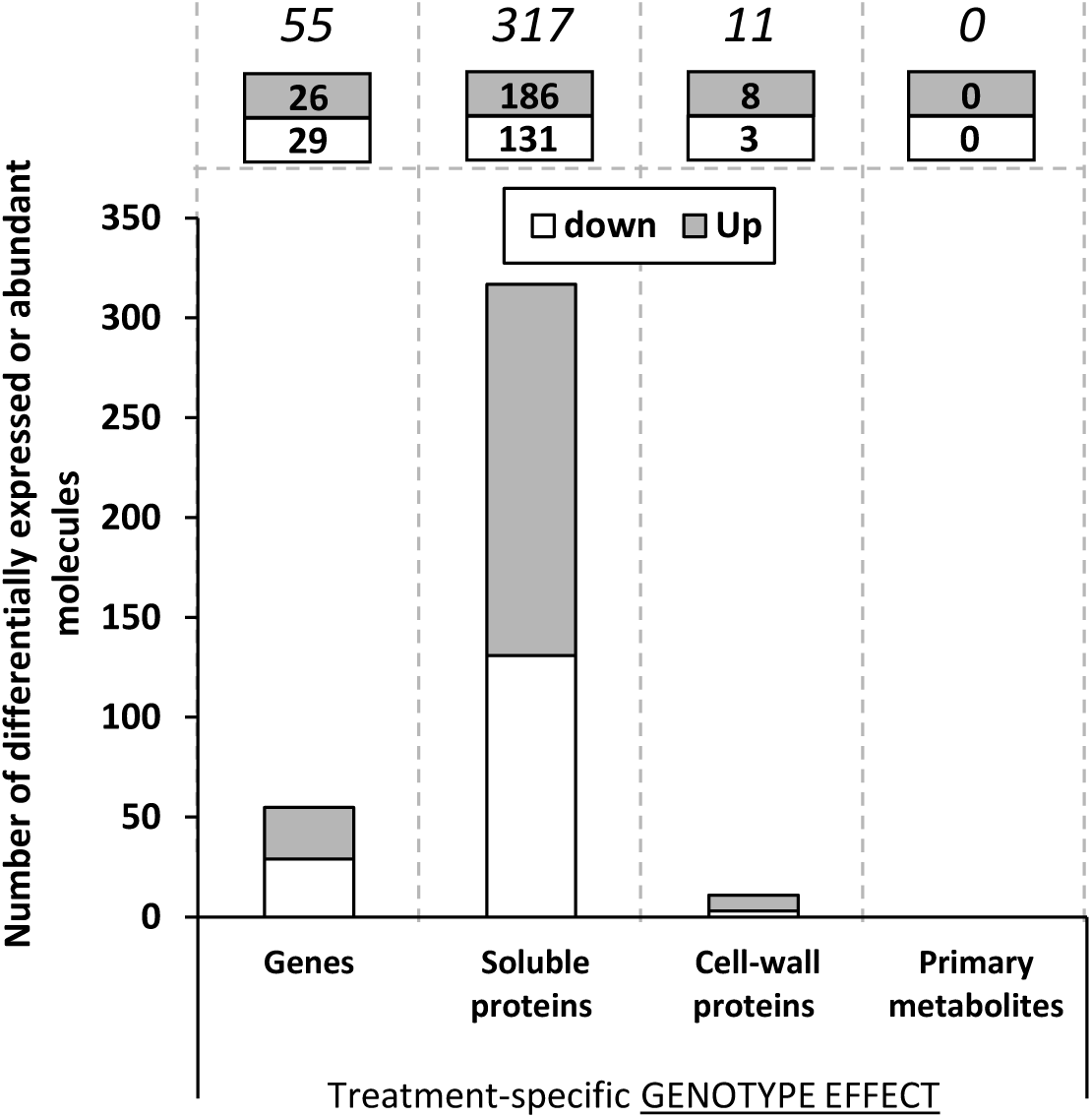
Overview of the number of genes, soluble proteins, cell-wall proteins and primary metabolites whose expression or abundancy were significantly modulated according to A) the genotype effect (gray bars, up; white bars, down). *The presence or absence of dots within bars indicates highest or lowest representation of the biological molecules within the panel, respectively. The analyzes were performed on maize leaves of M1 chill-resistant and M2 chill-sensitive genotypes subjected either to control or chilling treatment*.

**Fig. 5.**
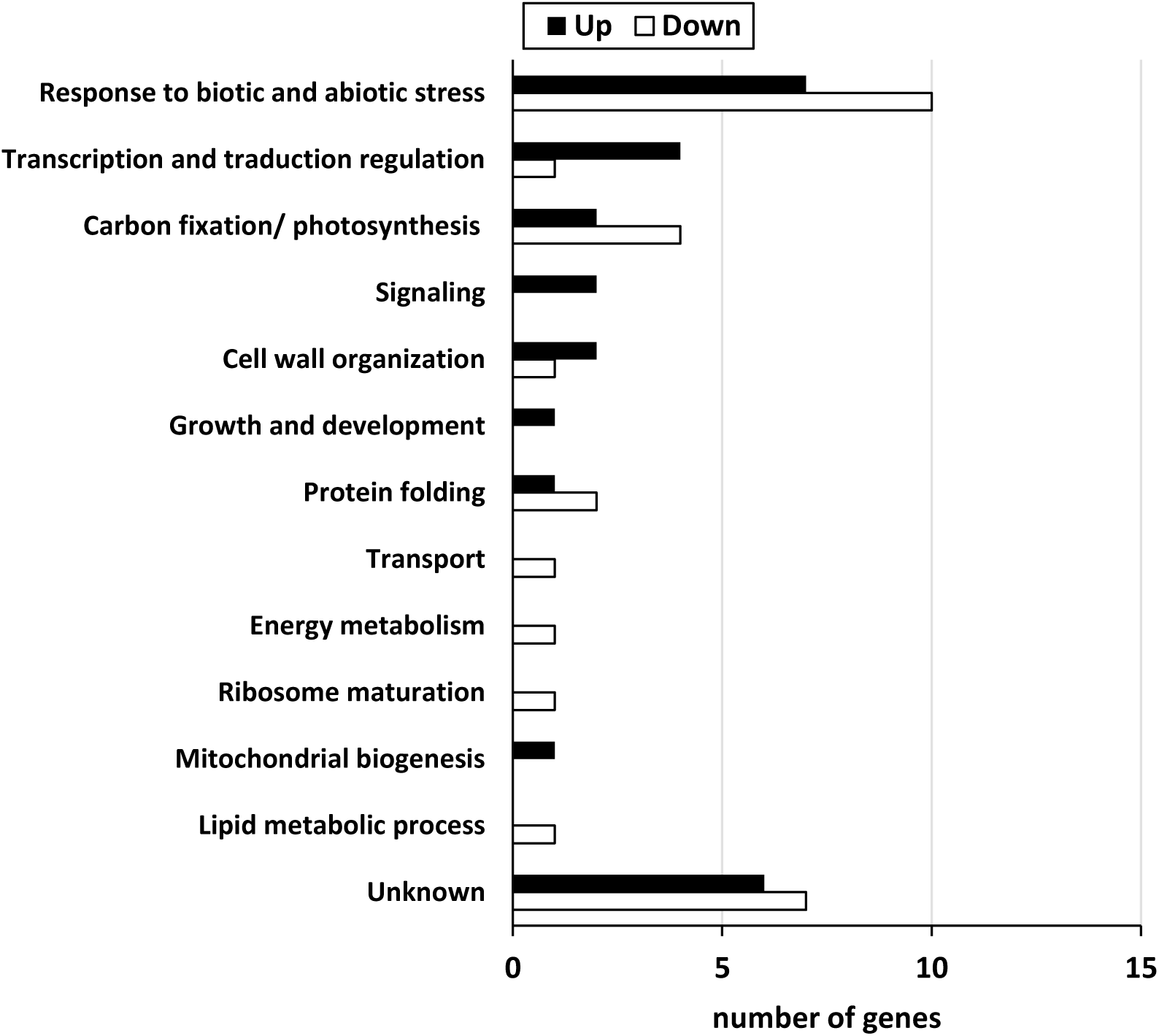
Biological functional annotations of differentially expressed genes (DEGs) linked to the genotype-effect by using Gene Ontology, Uniprot, KEGG, TAIR databases. *The black and white bars correspond to the up and down-regulated genes, respectively*.

### 3.4. Soluble proteome content

We observed a total of 317 DASPs with 131 and 186 more and less abundant soluble proteins (Figure 4; Table S1 and S2) which corresponded to 34 and 15 significantly enriched biological processes, respectively (Figure 6; Table S8 and S9). For DASPs with higher abundance in the M1 genotype compared to M2 during chilling treatment, there was a highly enriched biological process linked to the metabolic process (GO:0008152). Other biological processes that were less enriched in terms of protein number were involved in stress response, such as immune responses against viruses (GO:0051607), immune effector process (GO:0002252), secondary metabolite catabolic process (GO:0090487) and toxin catabolic process (GO:0009407). When considering the most down-abundant proteins, there was a very high enrichment of biological processes involved in the stress response, such as processes related to the immune response (GO:0006955). Surprisingly, biological processes of root development and morphogenesis (GO:0048364 and GO:0010015) were highly significantly enriched even though the fact that the tissue studied here was leaf. With highly significant p-values but few annotated genes, processes related to cell growth and cell wall formation were also enriched, including negative regulation of unidimensional cell growth (GO:0051511), negative regulation of cellular component organization (GO:0051129), regulation of cell wall organization or biogenesis (GO:1903338) and regulation of cell wall pectin metabolic process (GO:1902066). At last, the process associated to lipid transport (GO:0006869) was also enriched with a large number of proteins.

**Fig. 6.**
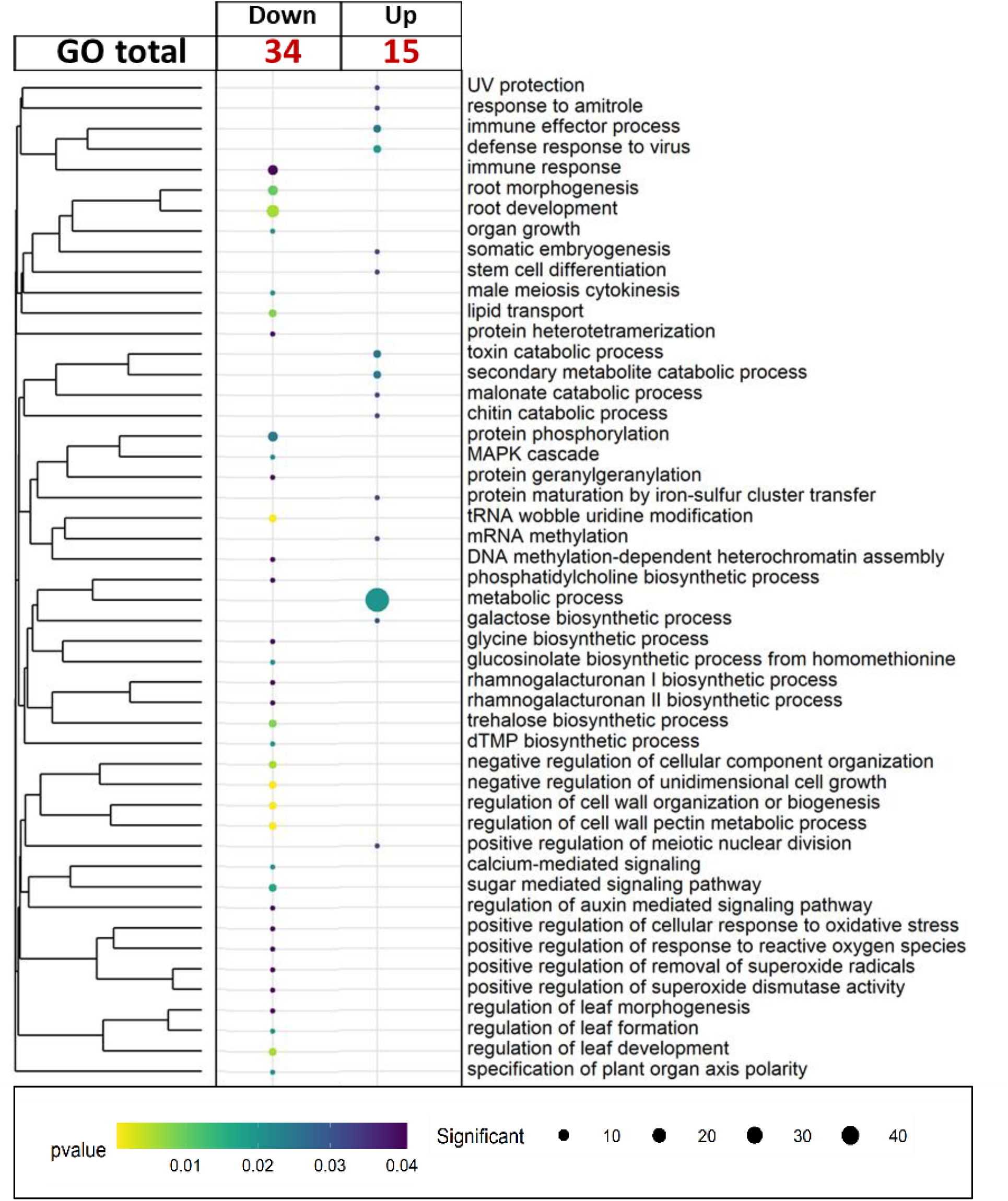
Bubble plot of enriched biological processes of more (up) or less (down) differentially abundant soluble proteins (DASPs) regarding to the genotype-effect. Biological processes are ordered by a semantic classification. *The size of the circle denotes the number of differentially abundant proteins (the biggest, the highest and the lightest the more significant, respectively). The color of the circles denotes the FDR (false discovery rate) adjusted p-value (the biggest the highest and the lightest the more significant, respectively). Biological processes enrichements were performed with topGO R package*.

### 3.5. Cell-wall proteome content

A total of 11 DACWPs was observed, with 8 more abundant proteins and 3 less in genotype M1 compared to M2 (Figure 4; Table 1; Table S3 and S4). Proteins associated with response to biotic and abiotic stresses showed the highest variations in abundance with logFC>5, such as glucan endo-13-beta-glucosidase (Zm00001d042143) and proteins responding specifically to oxidative stress such as a peroxidase (Zm00001d014606). Proteins whose abundance was the most reduced, with a LogFC<-5, are a laccase (Zm00001d023617) with potential roles in the stress response or in the cell wall organization and a dirigent protein (Zm00001d033942) with transport-related protein role. Proteins with less modulated abundances were involved in the response to stress, such as a Protein P21 (Zm00001d033455), a Beta-1,3-glucanase (Zm00001d042140) as well as proteins acting on the cell-wall organization such as Beta-expansin 1a (Zm00001d047096)

**Table 1.**
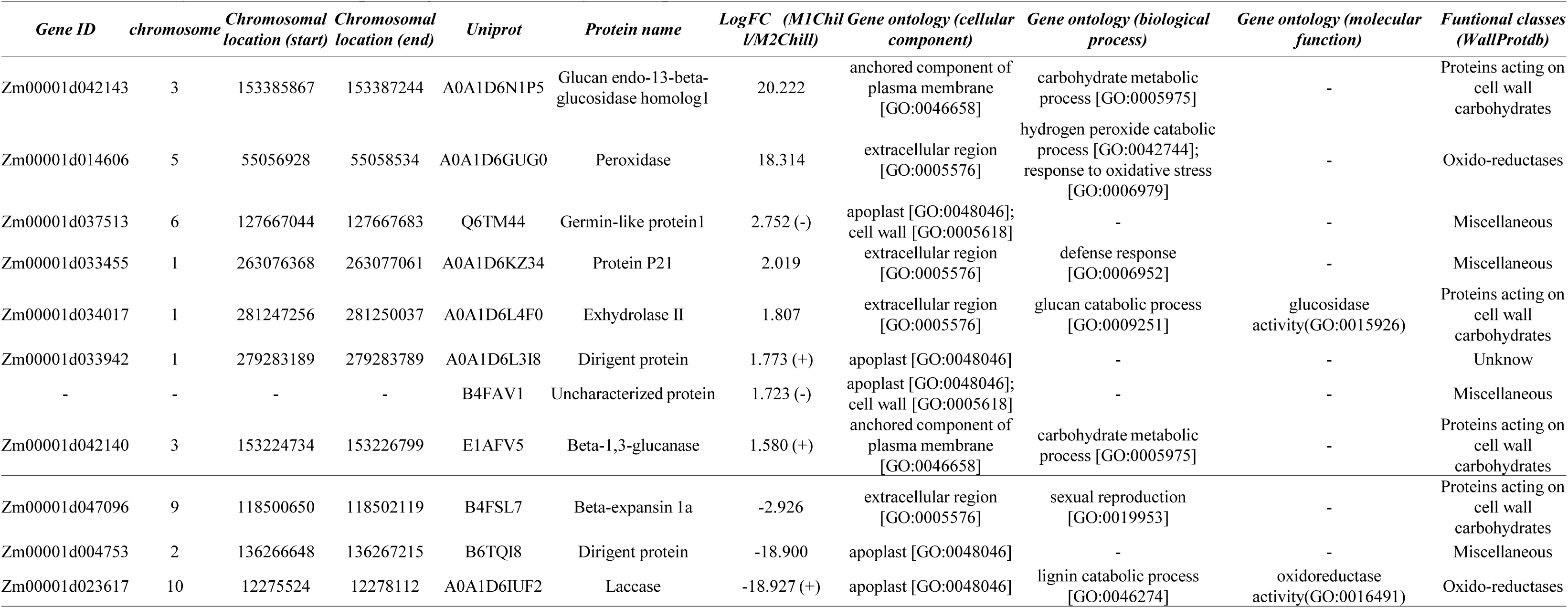
Significant differentially abundant cell-wall proteins (DACWPs) linked to the genotype-effect under chilling treatment. The functional classification was based on the functional classification of homologous proteins in WallProtdb (San Clemente and Jamet, 2015). The proteins whose response is constitutive under both conditions have a symbol (+) or (-) depending on whether they were up or down abundant.

### 3.6. Metabolome content

No primary metabolites exhibited differential abundance in response to the genotype effect (Figure 4). Among the 3569 metabolomic features analyzed, 61 variables showed significant differential abundance attributable to genotype (Table S7). When ranked by fold change, 9 of the 12 variables with the highest log₂ fold change (LogFC) were identified as benzoxazinoid metabolites, including BZX-like-glucoside, DIBOA-glucoside-2, and HBOA-glucoside-2 (Table 2).

**Table 2.** The twelve highest significant differentially abundant specialized metabolites linked to the genotype-effect under chilling treatment.

| *ID* | *LogFC*<br>(*M1Chill/M2Chill*) | *Adj. p-Value*<br>(*M1Chill/M2Chill*) | *Putative name* |
| --- | --- | --- | --- |
| X4627 | 2.471 | 0.0074 | BZXlike-glucoside |
| X4783 | 2.384 | 0.0218 | - |
| X2918 | 2.34 | 0.0188 | - |
| X2529 | 2.276 | 0.0074 | - |
| X1562 | 2.266 | 0.0074 | HBOA-Glucoside-2 |
| X2638 | 2.252 | 0.0076 | DIBOA-glucoside-2 fragment |
| X1585 | 2.243 | 0.0103 | DIBOA-glucoside-2 [M-H]- |
| X2552 | 2.236 | 0.0079 | DIBOA-glucoside-2 adduct |
| X2101 | 2.184 | 0.0074 | DIBOA-glucoside-2 adduct |
| X2085 | 2.176 | 0.0074 | DIBOA-glucoside-2 adduct |
| X201 | 2.15 | 0.0074 | DIBOA-glucoside-2 adduct |
| X1723 | 2.131 | 0.0074 | DIBOA-glucoside-2 adduct |

## 4. Discussion

Understanding the complexity of chilling adaptation requires integrating phenotypic, molecular and structural responses across scales and genotypes. In this study, morphometric measurements, physiological monitoring and a multilayered -omics strategy were combined in two near-isogenic maize lines (NILs) that differ only by a small genomic divergence on chromosome 4 (James et al., 2026). This design provides a powerful framework to dissect how a restricted genomic region can orchestrate extensive molecular rewiring and condition contrasting chilling tolerances. To our knowledge, this is the first work to consider transcriptomic, soluble-proteomic, cell wall–proteomic, metabolomic and physiological datasets to characterize chilling responses in maize NILs. The results reveal that a divergence affecting less than 0.15% of the genome can have major functional consequences and delineate pathways of interest for breeders.

### 4.1. Phenotypic plasticity: photochemical homeostatis and sustained leaf development

Chilling exposure induced clear and expected phenotypic modifications in both NILs, including reduced growth and declines in photosynthetic performance, consistent with previous observations in maize and other crops (Duran Garzon et al., 2019; Hajihashemi et al., 2018; Hassan et al., 2021; Ma et al., 2022; Tollenaar, 1989). The observed decreases in leaf length, visible leaf number and biomass reflected the well-known developmental inhibition caused by low temperatures. Reductions in chlorophyll content, Fv/Fm and ΦPSII confirmed impaired PSII function, in line with documented constraints on photosynthetic efficiency under chilling (Bilska-Kos et al., 2018; Riva-Roveda et al., 2016). These consistent treatment effects provided a robust foundation for interpreting subsequent genotype-dependent differences. Although shoot dry weight, leaf-3 length and width were unaffected, M2, the chilling-sensitive line, produced fewer total and ligulated leaves and exhibited significantly lower Fv/Fm than M1, the tolerant line. Given the unchanged dry weight, this pattern suggests that M2 leaves may be thicker under chilling, similar to the response observed in *Atriplex* (Riva-Roveda et al., 2016);. By contrast, both genotypes exhibited near-zero ΦPSII and chlorophyll content under chilling stress, indicating a strong inhibition of PSII activity and linear electron transport. Consequently, no genotype differences could be detected, consistent with reports of chilling-induced photosynthetic arrest in maize (Bilska and Sowiński, 2010; Foyer et al., 2002). The specific reduction in Fv/Fm in M2 therefore likely reflects enhanced susceptibility to photoinhibition. As shown in chrysanthemum under low temperature and varying irradiance (Janka et al., 2015), M1 may maintain or even increase PSII efficiency through superior thermal energy dissipation mechanisms. This enhanced photoprotection could arise from improved thermostability of the O₂-evolving complex or reaction centers, as reported in wheat (Lu and Zhang, 1999), or from modified photosystem stoichiometry, such as increased PSII relative to LHCII (Walters, 2005).

Beyond these phenotypic differences, chilling stress profoundly altered transcriptomic, proteomic, and metabolic profiles, as previously reported in maize (Li et al., 2019, 2017; Urrutia et al., 2021). Principal component analyses performed across the different omics layers confirmed a strong effect of the chilling treatment. In addition to this treatment effect, the multi-omics approach helped to clarify genotype-dependent responses. PCA of the soluble and cell wall proteomic datasets revealed clear genotype-specific patterns, allowing a robust discrimination between M1 and M2, and highlighted a large number of potentially key molecules that were differentially expressed or accumulated across the different omics layers. With the exception of primary metabolites, genotype-discriminating molecules identified in the other datasets were consistent with the phenotypic contrasts observed between the two genotypes (Figs. 3 and 4). Integrating the different omics datasets proved challenging due to the limited overlap between transcriptomic and proteomic data, with only a single common identifier detected: the O-methyltransferase ZRP4 (BX7). For this reason, each dataset was primarily analyzed independently, as they nevertheless revealed biologically meaningful and complementary key molecules. This conclusion was further supported by the multiple factor analysis (MFA), which reinforced the relevance of treating each omics layer separately. MFA integrating all datasets into a common multidimensional space (de Tayrac et al., 2009) showed that chilling stress accounted for the majority of the total variance (52%) in a consistent manner across all layers, whereas genotype explained 11% of the variance. The soluble proteome contributed the largest proportion of genotype-associated variance (approximately 60%), followed by the cell wall proteome (approximately 25%). Altogether, these results confirm the global impact of chilling stress on transcriptomic, proteomic, and metabolic responses in maize and demonstrate, for the first time, that the cell wall proteome undergoes extensive remodeling under chilling conditions.

### 4.2. Developmental responses: cell wall remodeling

Although such remodeling has not previously been documented in maize, it is consistent with the central role of the cell wall in response to chilling (Bilska-Kos et al., 2017; Le Gall et al., 2015). Cell wall proteins, although historically overlooked, are particularly relevant to chilling tolerance because the cell wall is the primary interface with environmental stress. Cold stress is known to alter cell wall proteins such as AGPs, extensins and XTHs (Le Gall et al., 2015) and to cause structural adjustments, including pectin and lignin accumulation, in frost-tolerant *Miscanthus* genotypes (Domon et al., 2013). In the present study, several hydrolases-Glucan like endo-1,3-β-glucosidase homolog 1, Exhydrolase II and β-1,3-glucanase-were more abundant in M1, while β-expansin 1a was reduced, indicating active remodeling of wall structure. Sun et al., (2025) previously reported that Glucan endo-1,3-β-glucosidase homolog 1 and Exhydrolase II are cold-induced and further enhanced under EBR treatment, consistent with differential brassinosteroid signaling pathway activity reported in the divergence region (James et al., 2026). Together with the differential abundance of resistance-related proteins and the cell-wall protein P21, these hydrolases are likely to contribute to the enhanced cold tolerance of M1. Glucan endo-1,3-β-glucosidase homolog 1 was particularly overaccumulated (LogFC = 20), underscoring its potential importance. Differences in apoplastic oxidative enzymes further distinguished the genotypes. A peroxidase (Zm00001d014606) was highly accumulated in M1 (LogFC = 18), whereas a laccase (Zm00001d023617) showed the opposite pattern (LogFC = −19). Cell-wall peroxidases are known to increase during cold stress alongside ROS accumulation (Sobkowiak et al., 2016; Theocharis et al., 2012). Cold-tolerant maize varieties typically maintain lower lipid peroxidation (Hu et al., 2025), and ascorbate peroxidase activity is higher in tolerant compared with sensitive genotypes (Pinhero et al., 1997). These observations strengthen the hypothesis that the identified peroxidase contributes to improved oxidative homeostasis and chilling tolerance in M1.

### 4.3. Stress and defense responses: metabolites and benzoxazinoids

Genotype-specific responses involved extensive regulation of biological processes related to stress, development, signaling, gene regulation and other core functions, in agreement with the broad physiological impact of cold (Ma et al., 2022; Waititu et al., 2021). In maize, very recent studies have shown that variation in brassinosteroid signaling components and downstream transcription factors contributes to in cold tolerance, consistent with broad effects on hormone signaling cascades and cold-responsive gene (Sun et al., 2025; Yang et al., 2025; C. Zhang et al., 2025). Numerous defense-related genes, including RPP13-like protein 4, RPM1 and RGA2, were strongly up-regulated in the tolerant genotype M1, particularly within the divergence region. These genes belong to families known to be modulated during cold stress (Wu et al., 2019) and may contribute to cross-protection mechanisms, given that homologs participate in drought or heat resilience (Li et al., 2024; Yang et al., 2021). In addition to these transcriptional changes, multiple non-targeted (RNA-seq and soluble proteomes) and targeted (specialized metabolites) approaches consistently pointed to the benzoxazinoid (BX) pathway as a central metabolic hub of the early chilling response. Within the divergent region, several benzoxazinoid biosynthetic genes (BX1, BX5, BX7) were down-regulated under chilling in the tolerant genotype M1, whereas BX6 was up-regulated. Although BX7 was the only gene showing a significant proteomic change, the tolerant line M1 accumulated the benzoxazinoids HBOA and DIBOA-glucoside (Figure 7). Targeted analyses of specialized metabolites showed that benzoxazinoid-related compounds represented around 16% of the 61 differentially abundant metabolites, confirming the strong focus on this pathway. This pattern is consistent with transcriptomic evidence indicating that benzoxazinoid pathways are enriched among genes responsive to drought and cold (Li et al., 2017) and with reports of benzoxazinoid induction under chilling in field conditions (Urrutia et al., 2021). Together, these observations support a model in which benzoxazinoids contribute to chilling tolerance in maize through roles in stress signaling, defense and modulation of oxidative balance. In this framework, the benzoxazinoid pathway appears as one of the main axes through which a small genomic divergence in a near-isogenic background translates into distinct metabolic and physiological outcomes.

**Fig. 7.**
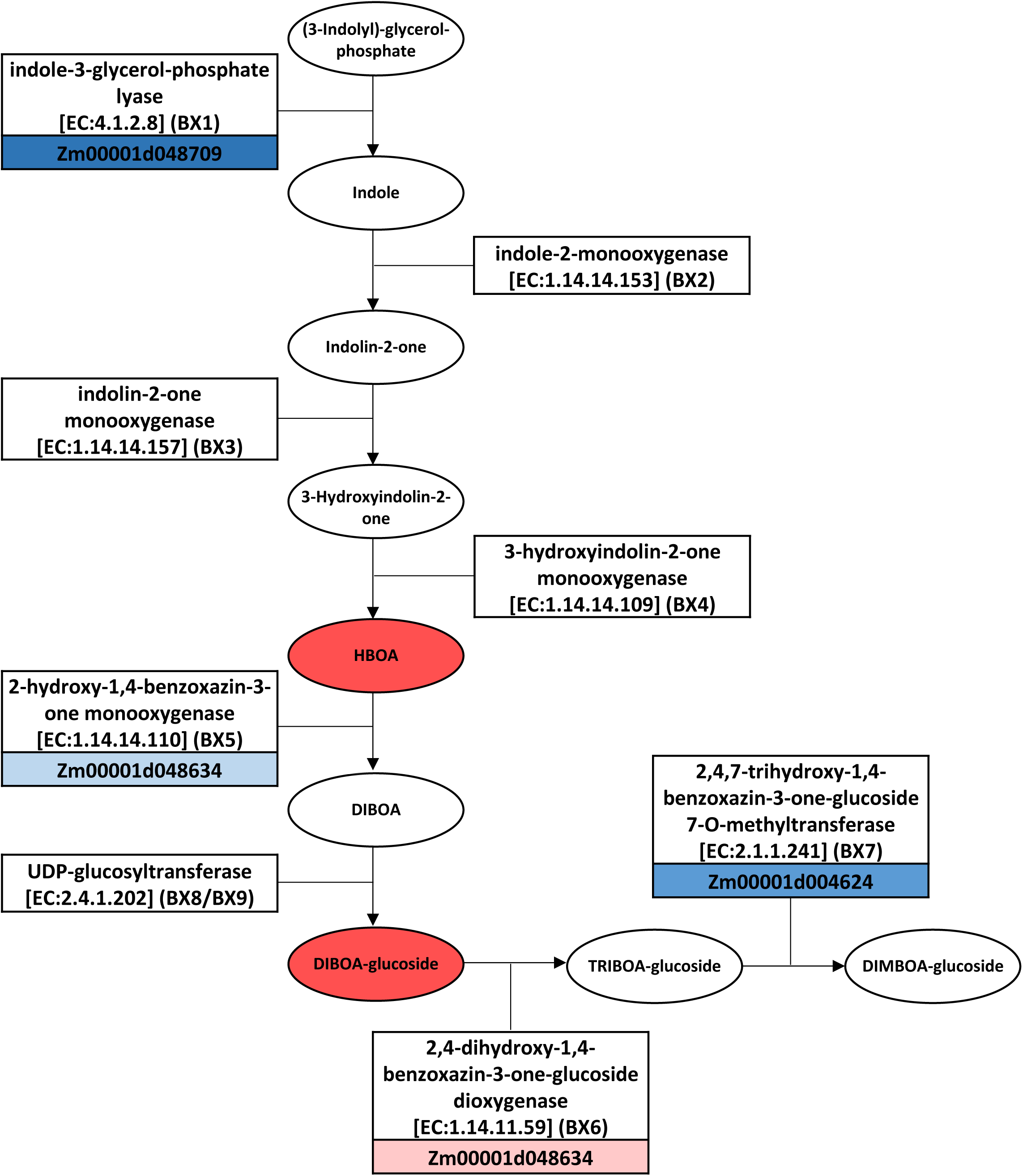
Representation of the known Benzoxazinoids biosynthetic pathway in *zea mays* L. and the regulation of the proteins and metabolites regarding to the genotype-effect. *The intensity of the red and blue colors are correlated with the LogFc of the overabundant and down-abundant proteins or metabolites. Proteins and metabolites involved in the pathways are from the KEGG database*.

Cold exposure disrupts cellular homeostasis and promotes a strong accumulation of reactive oxygen species (ROS) in plant cells. When produced in excess, ROS damage cellular membranes, notably through lipid peroxidation processes (Nadarajah, 2020). To cope with this imbalance, plants activate a wide range of protective and detoxification mechanisms. Oxidative stress responses are therefore considered a central component of plant adaptation to cold conditions (Zhou et al., 2022). The pronounced accumulation of a cell wall–localized peroxidase, together with increased levels of benzoxazinoids, both proteins and metabolites known to contribute to redox homeostasis, may explain the enhanced cold tolerance of the M1 genotype. In addition, esculetin, which was also found overaccumulated in the resistant genotype M1 (Table S5), is a coumarin reported to possess antioxidant and ROS-scavenging properties *in vitro* (Kim et al., 2008). This enhanced antioxidant capacity is also consistent with its improved photoinhibition response, reflected by higher Fv/Fm values in the resistant genotype. Furthermore, the strong induction and high abundance of numerous glutathione transferases involved in glutathione metabolism, observed at both the transcriptomic and soluble proteomic levels in genotype M1, support the hypothesis of a more efficient management of oxidative stress in this genotype compared with the sensitive genotype M2.

## 5. Conclusion **&** Perspectives

This work shows that a very small genomic divergence can orchestrate large, multi-scale differences in maize chilling responses, and that multi-layer statistics can reveal its underlying regulatory axes. It also delivers quantitative knowledge directly usable to refine process-based crop models and guide future phenotyping strategies under variable climates. The concomitant analysis of morphometric, physiological, transcriptomic, proteomic and metabolomic data in near-isogenic maize lines demonstrates that a very limited genomic divergence on chromosome 4 (<0.15%, ca. 5.1 Mb) can drive major differences in both long-term developmental adjustment and short-term stress and defense responses under non-freezing chilling. Soluble and cell-wall proteomes contribute more strongly than the transcriptome to genotype-related variance, highlighting cell-wall remodeling enzymes, apoplastic peroxidases and benzoxazinoid-associated pathways as key components of cold tolerance. Together, the organ-resolved, multi-omics integration pinpoints two main axes of response: (i) developmental and structural remodeling, especially at the cell-wall level, and (ii) rapid activation of stress- and defense-related pathways, including specialized metabolism, thereby providing molecular markers directly relevant for QTL mapping and marker-assisted selection in breeding programs targeting improved chilling tolerance. Building on these findings, targeted proteomics could be employed to identify novel QTLs associated with stress-responsive proteins, as demonstrated in drought studies (Blein-Nicolas et al., 2020), while targeted metabolomics on benzoxazinoids may help sort genotypes and even enable phenomic selection strategies (Rincent et al., 2018). Such integrative, targeted omics approaches have the potential to bridge molecular knowledge and applied breeding for improved chilling tolerance in maize.

## Abbreviations

CWP: Cell Wall Protein
DACWPs: Differentially Abundant Cell Wall Proteins
DASPs: Differentially Abundant Soluble Proteins
DEG: Differentially Expressed Gene
MFA: Multiple Factor Analysis
NILs: Near Isogenic Lines
PCA: Principal Component Analysis
QTL: Quantitative Trait Loci
ROS: Reactive Oxygen Species
SP: Soluble Protein

## CRediT authorship contribution statement

**Maxence James**: Data curation, Formal analysis, Investigation, Methodology, Validation, Visualization, Writing – original draft, Writing – review and editing. **Andrea Rau**: Data curation, Formal analysis, Investigation, Methodology, Writing – review and editing. **Anca Lucau-Danila:** Conceptualization, Investigation, Data curation, Formal analysis, Methodology, Supervision, Validation, Writing – review and editing. **Jean-Michel Saliou:** Investigation, Data curation, Writing – review and editing. **Bertrand Gakiere:** Data curation, Writing – review and editing. **Caroline Mauve**: Investigation, Data curation, Writing – review and editing. **Ludivine Soubigou-Taconnat:**. **Alexandra Launay-Avon:** Investigation, Data curation, Writing – review and editing. **Christine Paysant**: Investigation, Data curation, Writing – review and editing **Stéphane Bernillon:** Investigation, Data curation, Writing – review and editing. **Pierre Pétriacq:** Investigation, Data curation, Writing – review and editing. **Catherine Giauffret:** Conceptualization, Data curation, Funding acquisition, Investigation, Methodology, Project administration, Supervision, Visualization, Writing – review and editing. **Estelle Goulas:** Conceptualization, Data curation, Formal analysis, Investigation, Methodology, Project administration, Supervision, Validation, Visualization, Writing – original draft, Writing – review and editing.

## Acknowledgements

The author acknowledges all participants of this study for their outstanding contributions.

## Data availability

All data used in this article is presented. All omics data will be made publicly available in appropriate repositories once the manuscript is accepted.

## Supporting information

**Table S1.** Significant differentially less-abundant soluble proteins (DASPs) linked to the genotype-effect. The soluble proteins whose response is constitutive under both treatments are in bold. Gene ontology (GO) annotations, protein class and families/subfamilies were obtained with the Panther database (http://www.pantherdb.org/, Mi et al., 2019).

**Table S2.** Significant differentially more-abundant soluble proteins (DASPs) linked to the genotype-effect. The soluble proteins whose response is constitutive under both treatments are in bold. Gene ontology (GO) annotations, protein class and families/subfamilies were obtained with the Panther database (http://www.pantherdb.org/, Mi et al., 2019).

**Table S3.** Significant differentially less-abundant cell-wall proteins (DACWPs) linked to the genotype-effect. The cell-wall proteins whose response is constitutive under both treatments are in bold. The functional classification was based on the functional classification of homologous proteins in WallProtdb (San Clemente and Jamet, 2015). Gene ontology (GO) annotations, protein class and families/subfamilies were obtained with the Panther database (http://www.pantherdb.org/, Mi et al., 2019).

**Table S4.** Significant differentially more-abundant cell-wall proteins (DACWPs) linked to the genotype-effect. The cell-wall proteins whose response is constitutive under both treatments are in bold. The functional classification was based on the functional classification of homologous proteins in WallProtdb (San Clemente and Jamet, 2015). Gene ontology (GO) annotations, protein class and families/subfamilies were obtained with the Panther database (http://www.pantherdb.org/, Mi et al., 2019).

**Table S5.** Significant differentially less-expressed genes (DEGs) linked to the genotype-effect. The genes whose response is constitutive under both treatments are in bold. Gene ontology (GO) annotations, protein class and families/subfamilies were obtained with the Panther database (http://www.pantherdb.org/, Mi et al., 2019).

**Table S6.** Significant differentially more-expressed genes (DEGs) linked to the genotype-effect. The genes whose response is constitutive under both treatments are in bold. Gene ontology (GO) annotations, protein class and families/subfamilies were obtained with the Panther database (http://www.pantherdb.org/, Mi et al., 2019).

**Table S7.** Significant differentially abundant specialized metabolites linked to the genotype-effect.

**Table S8.** Biological processes enrichments of differentially more-abundant soluble proteins (DASPs). GO enrichment analyses were performed using TopGO with the *Zea mays* L. genome (B73_RefGen_v4) as a background.

**Table S9.** Biological processes enrichments differentially less-abundant soluble proteins (DASPs). GO enrichment analyses were performed using TopGO with the *Zea mays* L. genome (B73_RefGen_v4) as a background.

**Supplemental DataSet S1.** Total RNA-seq read counts in the genotypes M1 and M2 under control and chilling treatments.

**Supplemental dataset S2.** Raw Soluble protein data in the genotypes M1 and M2 under control and chilling treatments.

**Supplemental dataset S3.** Raw cell wall protein data in the genotypes M1 and M2 under control and chilling treatments.

**Supplemental dataset S4.** Raw primary metabolite data in the genotypes M1 and M2 under control and chilling treatments.

**Supplemental dataset S5.** Raw specialized metabolite data in genotypes M1 and M2 under control and chilling treatments.

